# Characterization of the Entner-Douderoff Pathway in *Pseudomonas aeruginosa* Catheter-associated Urinary Tract Infections

**DOI:** 10.1101/2023.11.14.567044

**Authors:** Nour El Husseini, Solomon A. Mekonnen, Cherisse L. Hall, Stephanie J. Cole, Jared A. Carter, Ashton T. Belew, Najib El-Sayed, Vincent T. Lee

## Abstract

*Pseudomonas aeruginosa* is an opportunistic nosocomial pathogen responsible for catheter-associated urinary tract infections (CAUTI). In a murine model of *P. aeruginosa* CAUTI, we previously demonstrated that urea within urine suppresses quorum sensing and induces the Entner-Douderoff (E-D) pathway. The E-D pathway consists of the genes *zwf, pgl, edd*, and *eda*. Zwf and Pgl convert glucose-6-phosphate into 6-phosphogluconate. Edd hydrolyzes 6-phosphogluconate to 2-keto-3-deoxy-6-phosphogluconate (KDPG). Finally, Eda cleaves KDPG to glyceraldehyde-3-phosphate and pyruvate, which enters the citric acid cycle. Here, we generated in-frame E-D mutants in strain PA14 and assessed their growth phenotypes on chemically defined media. These E-D mutants have a growth defect when grown on glucose or gluconate as sole carbon source which are similar to results previously reported for PAO1 mutants lacking E-D genes. RNA-sequencing following short exposure to urine revealed minimal gene regulation differences compared to the wild type. In a murine CAUTI model, virulence testing of E-D mutants revealed that two mutants lacking *zwf* and *pgl* showed minor fitness defects. Infection with the Δ*pgl* strain exhibited a 20% increase in host survival, and the Δ*zwf* strain displayed decreased colonization of the catheter and kidneys. Consequently, our findings suggest that the E-D pathway in *P. aeruginosa* is dispensable in this model of CAUTI.

**Importance:** Prior studies have shown that the Entner-Douderoff pathway is up-regulated when *Pseudomonas aeruginosa* is grown in urine. Pseudomonads use the Entner-Douderoff pathway to metabolize glucose instead of glycolysis which led us to ask whether this pathway is required for urinary tract infection. Here, single-deletion mutants of each gene in the pathway were tested for growth on chemically defined media with single-carbon sources as well as complex media. The effect of each mutant on global gene expression in laboratory media and urine was characterized. The virulence of these mutants in a murine model of catheter-associated urinary tract infection revealed that these mutants had similar levels of colonization indicating that glucose is not the primary carbon source utilized in the urinary tract.

## Introduction

Catheter-associated urinary tract infection (CAUTI) is the most common healthcare-associated infection, affecting approximately 15-25% of hospitalized patients who are catheterized during their stay (1). The primary risk factor for CAUTI is the increased duration of an indwelling catheter, with a 3-7% daily risk of developing an infection (1). This condition leads to extended hospital stays, with an annual cost ranging from 340 to 450 million, resulting from nearly 1 million cases each year (2). *Pseudomonas aeruginosa* contributes to 10% of CAUTIs and ranks among the top five most common pathogens associated with this condition (3). Unlike other causes of CAUTI, *P. aeruginosa* is not typically found as a part of the healthy human microbiome and is not considered a commensal organism (4). This unique characteristic makes *P. aeruginosa* a good candidate for the study of CAUTI since it eliminates the chance of recurring infection from a resident of the commensal microbiota. Studies on *P. aeruginosa* pathogenesis in CAUTI used a murine model based on a rat model (5, 6). In brief, the procedure involves using a catheter for transurethral deposition of an implant followed by the delivery of the inoculum into the bladder. In this murine model of CAUTI caused by *P. aeruginosa*, the strain PA14 exhibits both an acute phase and a chronic phase (7). The acute phase is characterized by weight loss, sepsis, and mortality, primarily driven by the type III secretion system (T3SS) (7). In contrast, the chronic phase is marked by weight gain and chronic colonization within the urinary tract, occurring independently of the T3SS (7). In a separate study, mutants of *P. aeruginosa* lacking *pel* or *psl* genes were present in similar numbers in the bladder and catheter as the wild-type bacteria during the CAUTI model, indicating Pel or Psl polysaccharides are dispensable for colonization in vivo (5). Bacterial RNA-sequencing of *P. aeruginosa* grown in urine or instilled into the mouse bladder revealed that quorum sensing is highly down-regulated (8). Testing mutants deficient in quorum signaling pathways, including Δ*lasR* and Δ*rhlRI*, demonstrated that quorum signaling is dispensable during murine CAUTI. Interestingly, this study also showed that *eda, edd, pgl*, and *zwf* genes were up-regulated (**Figure 1**) in urine and bladder instilled conditions when compared to phosphate-buffered solution with tryptone (PBS-T). This prompted us to hypothesize that the Entner-Douderoff (E-D) pathway may be important in CAUTI.

**Figure 1:**
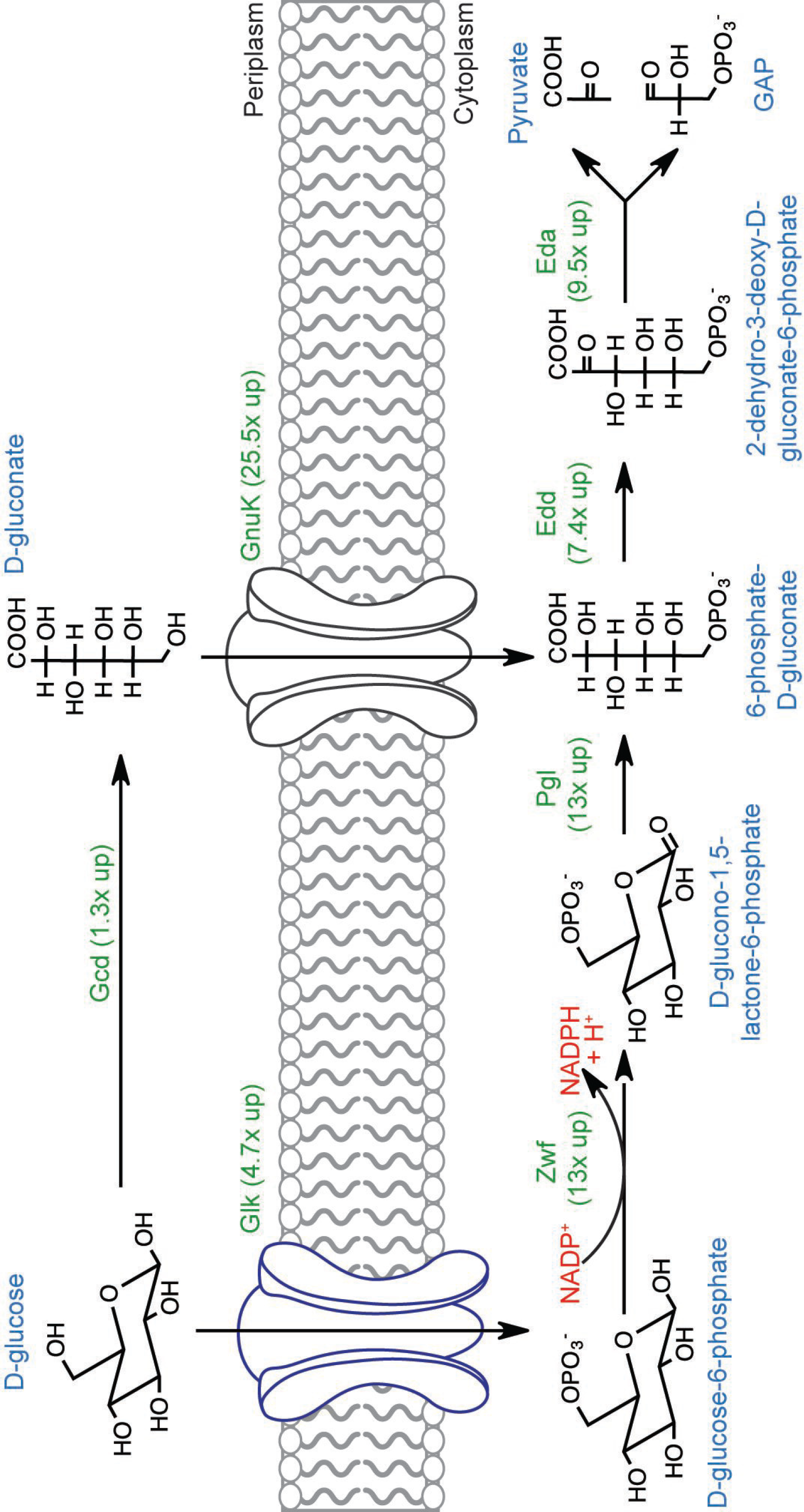
Genes encoding the enzymes for the Entner-Doudoroff (E-D) metabolic pathway are up-regulated during growth in urine and media with urea. Structures of the chemical intermediates of the E-D pathway are shown with names below the structure. The function of each enzyme is indicated above the arrow in green. The increased expression of each gene is indicated in parenthesis under the enzyme name in green as determined from Cole et al (8). Zwf oxidizes glucose-6-phosphate by reducing NADP+ to NADPH and H+ (indicated in red text).

Pseudomonads use the E-D pathway to convert glucose to glyceraldehyde-3-phosphate (GAP) and pyruvate because they lack the gene encoding phosphofructokinase-1, an enzyme that converts fructose-6-phosphate to fructose 1,6 bisphosphate in the classical glycolysis pathway (9, 10). Once glucose is imported into the periplasm by OprB (11), it can be converted to 6-phosphogluconate through oxidation and phosphorylation. This can occur through two routes referred to as the extracellular direct oxidative route and the intracellular phosphorylative route (10). In the direct oxidative route, glucose oxidation by Gcd in the periplasm yields D-gluconate, which is then phosphorylated and transported across the inner membrane by GnuK to yield 6-phosphogluconate. In the intracellular phosphorylative route, glucose is phosphorylated and transported across the inner membrane by Glk. D-glucose-6-phosphate is then oxidized in tandem by Zwf and Pgl into 6-phosphogluconate, where it can either be shunted to the pentose-phosphate pathway or continue in the E-D pathway. Within the E-D pathway, Edd hydrolyzes 6-phosphogluconate to 2-keto-3-deoxy-6-phosphogluconate (KDPG) (9). Finally, Eda cleaves KDPG to GAP and pyruvate, which enter the citric acid cycle for further metabolism (12). This pathway has been characterized in the *P. aeruginosa* strain PAO1 by early studies screening genetic mutants for growth defects on defined media and biochemical assays (10). From these studies, the genes encoding the enzymes responsible for the pathway were identified.

The contribution to virulence by the E-D pathway using mutant strains has been studied in several species but not yet in *P. aeruginosa*. Disruption of the E-D pathway in *Streptococcus pneumoniae* in a chinchilla model significantly increased virulence and decreased survival (13). In *Vibrio cholerae*, the E-D pathway was reported to be required for colonization in a suckling mouse model (14). Disruption of the E-D pathway decreases the fitness of *Helicobacter pylori* in a gastritis model in mice by decreasing colonization (15). In *Escherichia coli*, a MG1655 *edd* mutant had a defect in colonization of the mouse gastrointestinal tract (16). Another study found an *eda* mutant in both the F-18 and K-12 strains was unable to colonize the intestines in Streptomycin-treated mice (17). In contrast, an *E. coli* CFT073 *edd* mutant colonized similarly to the parental strain in a murine UTI model (18). These studies demonstrate that the requirement of the E-D pathway varies among species and the site of infection. This knowledge gap, along with our previous RNA-seq results, motivated us to investigate the E-D pathway in *P. aeruginosa* CAUTI. Here, our objectives were to evaluate mutants of these genes in PA14 for their contribution to global transcriptional regulation and virulence in a CAUTI murine model.

## Results

### Growth of strain mutants on minimal media with single carbon sources

*P. aeruginosa* genetic mutants in the E-D pathway have been previously identified or generated in PAO1. To test the effect of these genes in the PA14 strain used previously in our established murine model of CAUTI, in-frame deletion mutants of each gene in the E-D pathway were created by homologous recombination and verified by sequencing in PA14. Each strain was tested in growth assays by spotting 10-fold serial dilutions on chemically defined media. When tested on plates with citrate as the sole carbon source, all strains grew equally (**Supplementary Figure 1**) as this metabolite can directly enter the citric acid cycle and bypass the upstream E-D pathway. When tested on plates with glucose as the sole carbon source, wild-type PA14 grew on glucose with the same plating efficiency as on citrate (**Figure 2**). However, Δ*eda* had a 6-log growth defect, Δ*edd* and Δ*pgl* strains had a 5-log growth defect, and Δ*zwf* had a 3-log defect. These results suggest each of the E-D genes is necessary for glucose metabolization and normal growth on glucose. In contrast, the Δ*gcd* mutant had no growth defect, suggesting that extracytoplasmic conversion of glucose to gluconate by Gcd is not essential for glucose metabolism. The growth defect on glucose was fully complemented by expression of the corresponding gene in trans from a plasmid for Δ*eda*, Δ*pgl*, and Δ*zwf* ; while expression of *edd* from a plasmid restored plating efficiency by 4-logs. However, expression of *gcd* from a plasmid reduced growth by 1-log. Overexpression of *gcd* or *edd* lead to a 6-log or 4-log reduction in plating efficiency, respectively, even on citrate (**Supplementary Figure 1**). These results suggest that over-expression of *gcd* or *edd* leads to toxic effects in *P. aeruginosa*.

**Figure 2:**
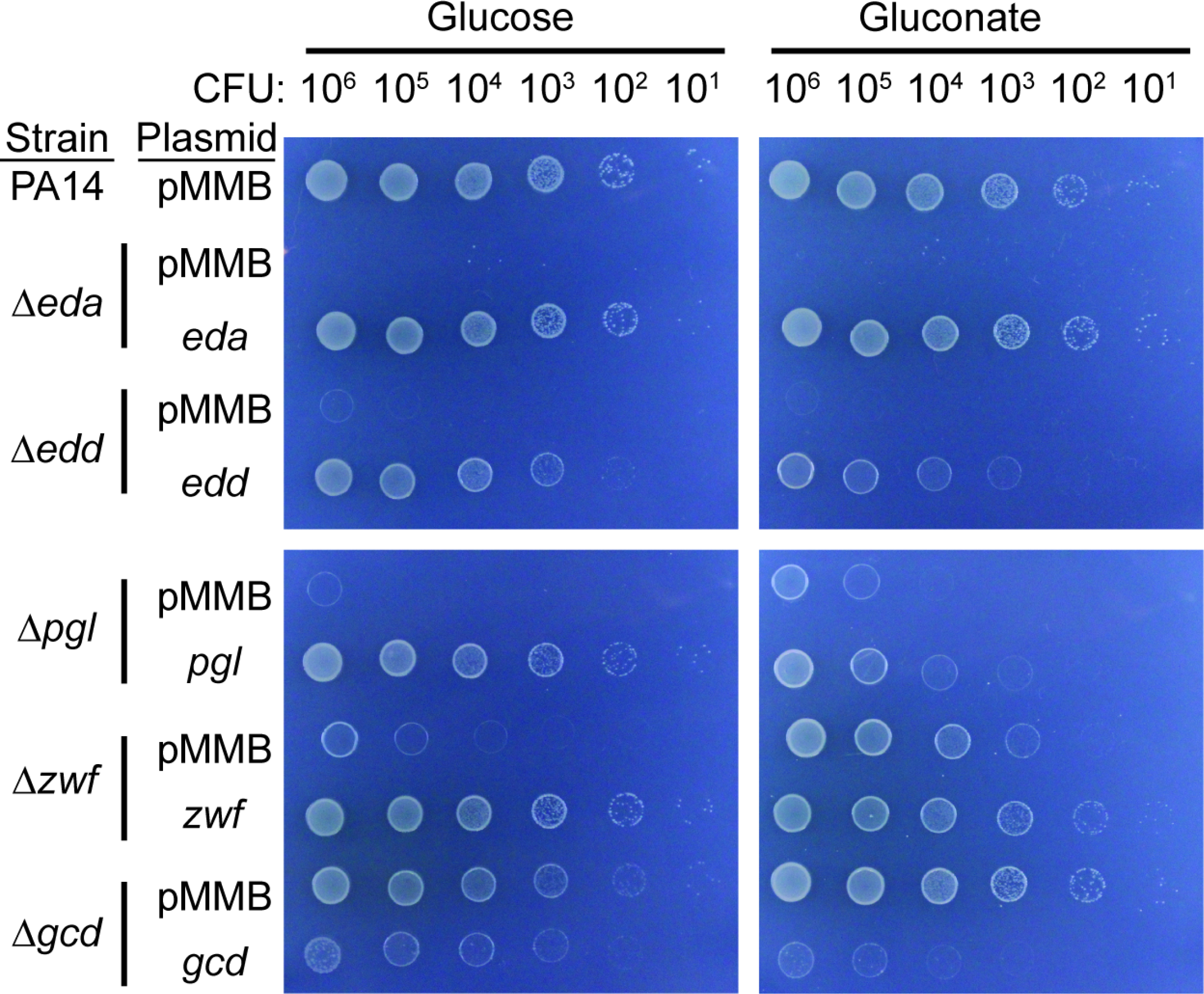
Requirement of E-D pathway genes for growth on glucose or gluconate as a sole carbon source. Indicated strains with specified plasmid were serial 10-fold diluted and 5 *µ*L were spotted on the plates with the specified carbon source and 1 mM IPTG to induce the *tac* promoter on the plasmid.

When tested on plates with gluconate as the sole carbon source, parental PA14 grew on gluconate with similar efficiency as glucose and citrate. The Δ*eda* and Δ*edd* mutants had a 6-log reduction in plating efficiency, suggesting they are both essential for gluconate metabolism. Δ*pgl* and Δ*zwf* mutants had attenuated growth of 4-log and 2-log defects, respectively. While these genes are upstream and do not play a direct role in gluconate metabolism, their decreased plating efficiency suggests a regulatory role or reduced efficiency of GnuK in translocating gluconate. The Δ*gcd* mutant grew similarly to the parental PA14, suggesting that Gcd is not essential for gluconate metabolism. The growth defect on gluconate of each of the E-D mutant was complemented by expression of the corresponding gene in trans from a plasmid. Similarly to glucose, the growth defect on gluconate was fully complemented in *eda* and *zwf* strains and partially complemented in *edd* and *pgl*. Overexpression of *gcd* led to a similar 3-log reduction in plating deficiency. Altogether, these results suggest PA14 metabolizes glucose through two routes, direct oxidation through Gcd and intracellular phosphorylation through Zwf and Pgl to produce the intermediate metabolite, and confirm that Edd and Eda are essential for complete metabolism in a manner similar to PAO1.

### Growth of E-D mutants on complex media sources containing urea

Previously, E-D pathway genes were shown to be highly up-regulated when *P. aeruginosa* is grown in urine and urea (8). The requirement of E-D pathway genes for growth on 75% human urine was tested using mutants harboring the corresponding genes from a plasmid that was induced with IPTG (**Figure 3**). PA14 grew normally on human urine. The Δ*eda* and Δ*edd* mutants showed minor plating defects of 1-log and 2-log decreases, respectively. The Δ*pgl*, Δ*zwf*, and Δ*gcd* mutants grew similarly to the parental PA14 strain. Complementation of *edd* or *eda* mutant strains with the corresponding deleted gene restored growth to PA14. Interestingly, the same growth defect was not observed upon overexpression of *edd* and *gcd* as seen in citrate, glucose, and gluconate. The requirement of E-D pathway genes was then tested for growth on lysogeny broth (LB) supplemented with 0.5% urea (**Supplementary Figure 2**). All strains grew nearly equally on urea. These results suggest *P. aeruginosa* primarily use metabolic pathways other than E-D pathway when other complex carbon sources are available in the growth media.

**Figure 3:**
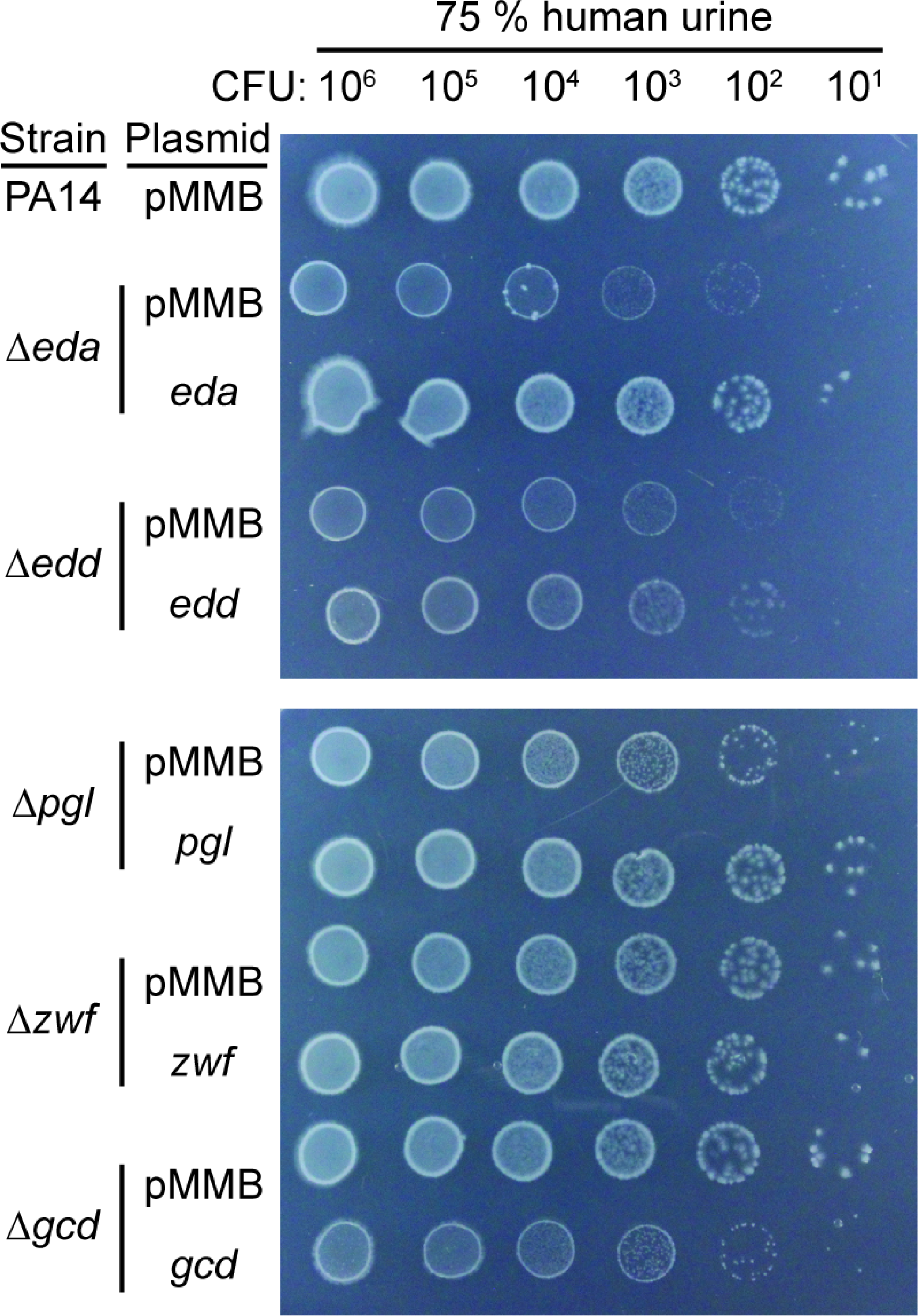
Requirement of E-D pathway genes for growth on pooled human urine. Indicated strains with specified plasmid were serial 10-fold diluted and 5 *µ*L were spotted on the plates with 75% human urine and 1 mM IPTG to induce the *tac* promoter on the plasmid.

### RNA-seq of mutant strains

Urea induces transcriptional changes in *P. aeruginosa*, including up-regulation of the glucose metabolism genes (8). To determine if these genes created conditions that regulated global expression, we tested the mutant strains in CAUTI using RNA-sequencing. Global expression in the absence of each gene was measured in vitro and in vivo. To measure transcriptional changes to urine in absence of a host immune response, we grew each strain in mouse urine and compared to their transcriptomes to the bacteria in the inoculum used for the CAUTI experiment (5). To measure transcriptional changes to urine and host immune responses, strains were instilled in a mouse bladder to mimic CAUTI. After one hour, contents of the bladder were collected, preserved in RNAlater and processed. In vitro incubation of samples in mouse urine and PBS-T was limited to two hours to approximate the time scale for the in vivo bladder samples. Total RNA was isolated, depleted of ribosomal RNA, and Illumina sequenced. The samples from bacteria grown in vitro yielded 2.9 million to 6.8 million reads mapped to the *P. aeruginosa* PA14 reference genome per sample. The samples isolated from the bladders of mice yielded 3.5 million to 5.9 million reads mapped to the *P. aeruginosa* and the rest mapped to the *Mus musculus* reference genome. The mouse transcriptome had an average of 4.3 million reads per sample and was not analyzed due to insufficient read depth.

To determine if resuspension of the inoculum in PBS-T changes the transcriptome of bacteria, samples were clustered to determine effects of strain and media using principal component analysis (PCA) on variance-stabilized transformed reads (19). We plotted samples grown in LB and PBS-T (**Supplementary Figure 3**). LB samples clustered together with PBS-T samples, suggesting that bacteria grown in LB and resuspended in PBS-T have a similar bacterial transcriptome. Since the transcriptomic signature of LB and PBS-T are largely similar, we used the PBS-T samples to compare samples incubated in mouse urine and instilled in mouse bladder. Principle component (PC) analysis revealed three distinct clusters corresponding to each condition, that is incubation in PBS-T, mouse urine, or instillation into the mouse bladder (**Figure 4A**). PC 1 contributed to 58% of the variance and differentiated mouse instilled samples apart from in vitro conditions PBS-T and mouse urine. PC 2 contributed to 25% of the variance which differentiated between the in vitro conditions PBS-T and mouse urine. These findings suggest the main drivers of different gene expression were due to the condition rather than genetic differences in the different strains. Since the clustering of samples through PCA suggested the mutants are similar to the parental PA14 in each condition, we verified expression of the E-D genes in these mutant strains. While PA14 strain samples expressed all E-D genes, the expression of the corresponding mutant strain was absent in the region deleted while other E-D genes retain their expression (**Figure 4B**). Clustering analysis using only the expression of the E-D genes showed that the samples clustered due to strain instead of media condition.

**Figure 4:**
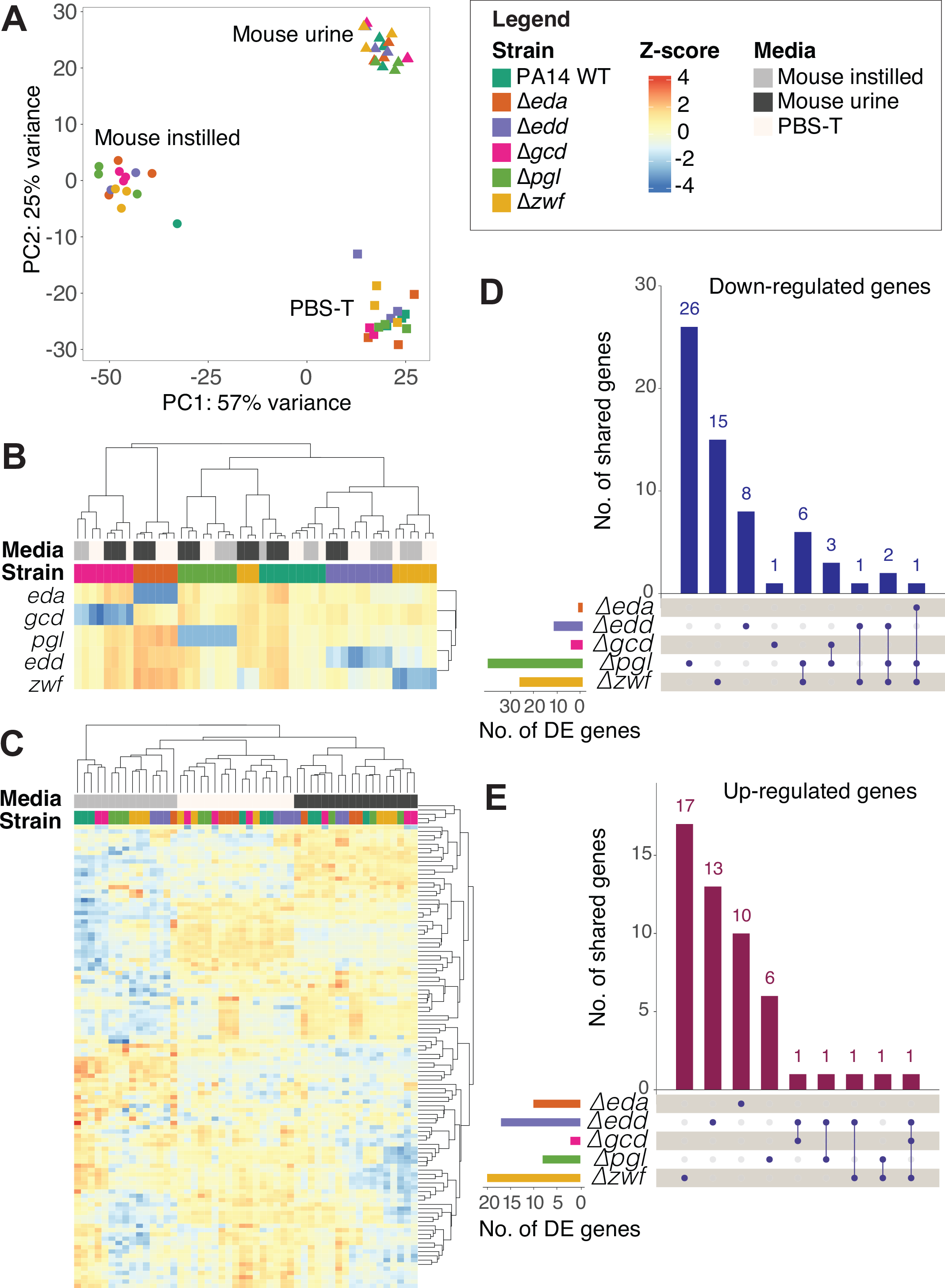
Global gene expression profiles of *P. aeruginosa* parental PA14 and E-D mutants in several growth conditions. Bacteria were incubated either PBS-T (non-induced growth) or mouse urine for two hours, or instilled in a mouse bladder for one hour. RNA was isolated, ribosomal RNA depleted, and sequenced. Subsequent analyses were performed using all *P. aeruginosa* annotated genes (5888) after filtering for low counts and variance-stabilizing normalization using DESeq2. **A)** Principal components analysis clustering of strain and media. Each point represents a biological sample with point color indicating *P. aeruginosa* strain and point shape indicating growth condition. **B)** log_2_ normalized gene expression heatmap of only E-D genes in all strains. **C)** log_2_ normalized gene expression heatmap of all differentially expressed (DE) genes comparing mutant strains against parental PA14 in each medium (≥ 2 or ≤−2 log_2_ fold change and a ≤ 0.05 *P* value cutoff). UpSet plot displaying **D)** down-regulated and **E)** up-regulated DE genes across mutant strains in all conditions. Sets on the left represent the total number of DE genes for each strain, indicated by horizontal bars. Intersections of shared DE genes are denoted as dots within the matrix, with the size of each intersection indicated by the vertical bars.

To investigate the impact of the E-D pathway on gene expression, we conducted a differential expression analysis comparing the parental strain to each deletion mutant in every experimental condition (**Supp. Table 1**). In the PBS-T condition, we identified a total of 21 up-regulated genes (defined as ≥ 2log_2_ fold change) and 10 down-regulated genes (defined as ≤−2log_2_ fold change) across all mutants when compared to the parental strain, using a ≤ 0.05 adjusted *P* value cutoff. In the pooled mouse urine condition, we found 12 up-regulated genes and 9 down-regulated genes. In the mouse instilled condition, there were 43 up-regulated genes and 85 down-regulated genes.

To gain insights into the functions of these differentially expressed genes, we created a heatmap displaying all differentially expressed genes from each experimental condition (**Figure 4C**). The columns in the heatmap cluster together, primarily due to the experimental conditions, as shown in the dendrogram. The functions of the differentially expressed genes were predominantly related to energy metabolism, specifically electron transport, polysaccharide and fatty acid metabolism, amino acids, and pathways such as the pentose phosphate pathway and E-D pathway.

To assess the overlap of differentially expressed genes among coregulated E-D pathway mutants, we employed an UpSet plot to visualize shared down-regulated (**Figure 4D**) and up-regulated (**Figure 4E**) genes across all media conditions. Our expectation was that the coregulated *edd* and *eda* strains would exhibit similarity, as would the *pgl* and *zwf* strains. However, the majority of differentially expressed genes appear to be unique to specific strains, with Δ*pgl* and Δ*zwf* having 26 and 15 distinct down-regulated genes, respectively. Notably, there was no overlap in down-regulated genes between the Δ*eda* and Δ*edd* mutants, while the coregulated *pgl* and *zwf* mutants shared only 6 down-regulated genes. In terms of up-regulated genes, Δ*pgl* and Δ*zwf* mutants exhibit the highest numbers, with 13 and 17 genes, respectively. Only one gene is up-regulated in both Δ*eda* and Δ*edd*, as well as Δ*pgl* and Δ*zwf*. The limited number of shared genes between coregulated E-D pathway mutants can be attributed to the overall low number of differentially expressed genes compared to the parental strain. These results suggest that the genes of the E-D pathway do not directly regulate gene expression during these conditions.

### E-D mutants in murine CAUTI model

While the E-D pathway genes do not contribute to transcriptomic changes in the conditions tested, whether the genes provide a functional role during infection remained to be determined. To test this question, we used the murine model of CAUTI with outbred CF-1 mice as previously described (8). In brief, after one week of acclimation, several cohorts of mice were infected through a transurethral procedure with parental PA14 or one of the isogenic E-D mutants Δ*eda*, Δ*edd*, Δ*pgl*, or Δ*zwf*. Mice were monitored daily for clinical signs, body weight changes, and survival for 14 days. Mice were sacrificed at the first sign of morbidity or upon termination of the experiment on day 14. Following termination, mice were dissected and bacterial burdens in the catheter, bladder, and kidneys were quantified. Survival analysis (**Figure 5A**) showed that all strains had similar biphasic infection as previously reported for the parental PA14 (7). Higher mortality rates were observed during the first 7 days, and few mice succumbed to infection in the last 7 days. Compared to parental PA14, the Δ*pgl* mutant showed a slower rate of mortality resulting in an 20% increased survival, while the rest of the mutants did not show differences in survival. In the catheter, Δ*zwf* had decreased colonization over the PA14 strain, whereas the other mutants had similar levels to PA14. Colonization in the bladder revealed no differences in abundance between PA14 and the mutant strains (**Figure 5B**). Finally, the Δ*zwf* mutant had a lower bacterial burden than PA14 in the kidneys while the rest of the mutants colonized similarly to PA14. There are subtle differences for some of the mutants in some aspects of the infection, suggesting that the E-D pathway provides a minor contribution in the pathogenesis of *P. aeruginosa* CAUTI in healthy outbred mice.

**Figure 5:**
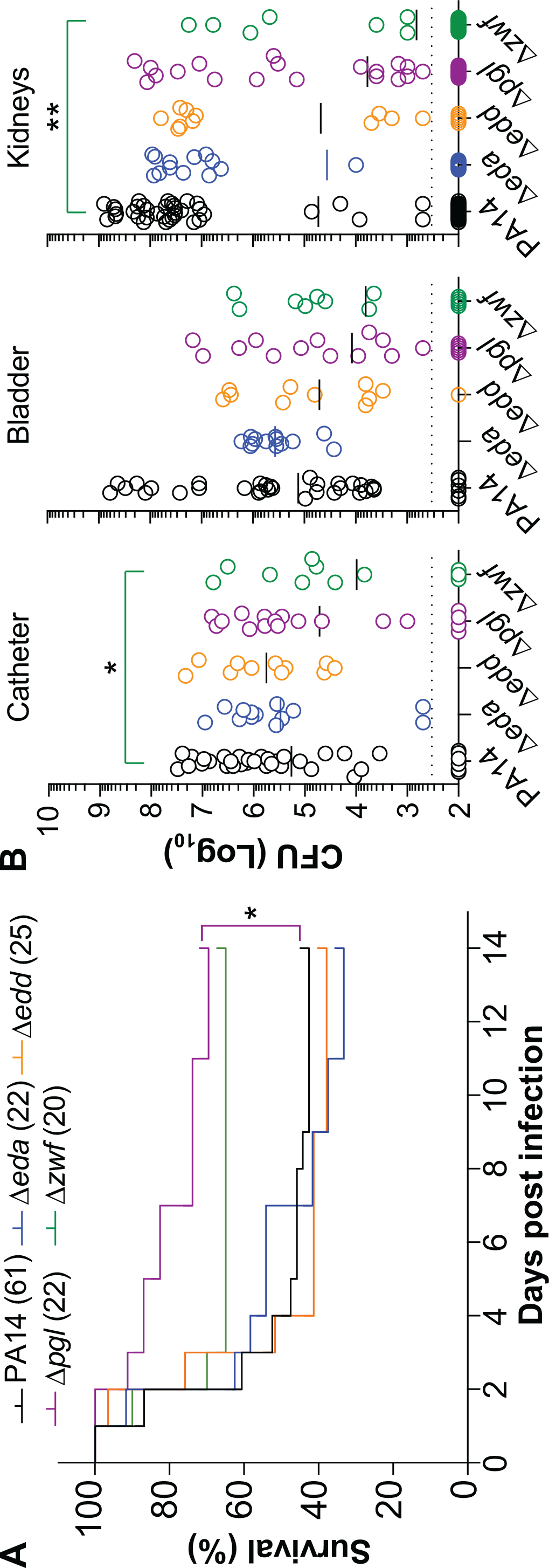
PA14 strains with mutations in E-D genes have minor defect in murine model of CAUTI. CF-1 transurethral infected with 3.5 × 10^7^ CFU of the indicated strain and monitored for morbidity and mortality for 14 days. **A)** Survival curve of mice infected with indicated strains. **B)** CFU burden of catheter, bladder, and kidneys with animals infected with the indicated strains at the time of sacrifice. PA14 data was previously published (7). Statistical analysis for survival curves between mice infected with PA14 and mutant strains was performed using a stratified log-rank test (Mantel–Cox). Statistical analysis for colony counts between mice infected with PA14 and mutant strains was performed using a Mann-Whitney U test. ^*^*P <* 0.05, ^**^*P <* 0.01

## Discussion

Growth assessment for the PA14 strains with in-frame deletion mutations generated in this study can be compared to mutants from early studies in the strain PAO1. These studies from the 1970s used chemical mutagenesis with *N*-methyl-*N*’-nitro-*N*-nitrosoguanidine, which can cause multiple mutations on the chromosome (20). To identify the location of these mutations responsible for metabolism of glucose and gluconate, mutants were tested for growth defects on single carbon sources. These mutants were characterized biochemically from cell lysates through enzymatic assays for the E-D pathway. Subsequent identification of spontaneous revertants that restored growth and biochemical activity validated their findings. PAO1 *edd* mutants (21–23) and *eda* mutants (23) did not grow in glucose or gluconate, confirming their essential role in metabolism. However, alternative routes can metabolize glucose to yield the 6-phosphogluconate intermediate, as evidenced by the *zwf* mutant having a minor defect in glucose (22, 24) and no defect in gluconate (22–24). These PAO1 mutants matched the expected phenotype based on the characterized metabolic pathway, suggesting that potential additional mutations from chemical mutagenesis did not affect growth. A later study generating a Δ*pgl* strain showed a 50% growth defect in gluconate as the main carbon source with minimal succinate and was complemented with a plasmid containing *pgl* (25). This result is intriguing since neither carbon source presumably requires Pgl for metabolism, suggesting a regulatory role for the gene. In the current study, we generated mutant strains in PA14 and tested them for growth on single carbon sources. As expected, all strains grew equally in citrate, which enters into the citric acid cycle downstream from the E-D pathway. Similar to PAO1, the PA14 Δ*eda* and Δ*edd* mutants did not grow in glucose or gluconate. The Δ*zwf* mutant had a major (≥ 3-log) plating defect in glucose and a minor (≤ 2-log) plating defect in gluconate, while the Δ*pgl* mutant had a major plating defect in both conditions. Defects in the latter two mutants are expected in glucose, but their defects in gluconate suggest a regulatory role similar to the Δ*pgl* strain in PAO1. Minor differences in phenotypes between PAO1 and PA14 observed in the *pgl* and *zwf* mutants could be due to differences in conditions; PAO1 strains were grown in liquid medium, and PA14 strains were grown on agar plates. We confirmed these results in PA14 through complementation of the corresponding genes, which restored growth to the mutants showing a growth defect. During these complementation studies, we observed that overexpression of *gcd* and *edd* led to a decrease in plating efficiency even when growing on citrate, a carbon source that bypasses the E-D pathway, suggesting that elevated levels of these two proteins have toxic effects in the cell. Our findings are also in support by a study using ^13^C-metabolic flux analysis to show increased intermediate in the E-D pathway when *P. aeruginosa* PAO1 was grown on glucose as sole carbon source (26). Their findings also extended to UTI and CAUTI isolates grown on glucose as a sole carbon source. Overall, these results indicate that these PA14 mutant strains have similar growth phenotypes on glucose and gluconate that corresponds to prior studies with PAO1 mutant strains (10).

Motivated by our recent finding that *P. aeruginosa* up-regulated the E-D pathway in mouse urine (8), we tested the growth of the PA14 mutant strains in complex-carbon media sources containing urea. All strains grew equally in LB supplemented with 0.5% urea. For this experiment, we used human urine because collection of mouse urine was limited.

When tested in 75% human urine, the Δ*edd* and Δ*eda* mutants had minor plating defects while the rest of the strains grew like WT. This shows the E-D pathway in *P. aeruginosa* contributes little to growth when other carbon sources are available. Furthermore, the glucose content is often low in urine of healthy individuals which could explain the moderate effects. This is one of the first reports examining the behavior of *P. aeruginosa* E-D mutants in media containing urea and urine. These results align with previous research highlighting the preference in *P. aeruginosa* for amino acids and organic acids, such succinate, pyruvate or acetate, as energy sources over glucose (27).

To investigate environmental adaptation changes through transcriptional responses, we conducted RNA-seq analysis on *P. aeruginosa* PA14 strains grown in mouse urine or instilled into the bladder and compared these results to the transcriptome of bacteria resuspended in the inoculation media. We did not observe the same E-D signature as in our previous study, where the E-D pathway was up-regulated in urine (8). In these current experiments, the bacteria were centrifuged, resuspended in new media, and incubated to better mimic the inoculum used for CAUTI experiments rather than subcultured and grown in media as in the prior study. These experiments should have allowed us to detect bacterial changes that are specific to host responses that is beyond exposure to urine. Our findings indicate approximately 20 differentially expressed genes (DEGs) (defined as ≥ 2 or ≤−2log_2_ fold change and ≤ 0.05 adjusted *P* value) between bacteria grown in LB and those resuspended in PBS-T (**Supplementary Figure 4**), suggesting that the preparation of the bacteria for inoculation has minimal impact on gene expression. However, in vitro incubation in filtered mouse urine had approximately 400 DEGs compared to PBS-T. This alteration became more pronounced in vivo, with bladder instillation in a mouse resulting in approximately 1100 DEGs (**Supp. Table 2**). To contextualize our results, we can compare them to a previous study that assessed the transcriptional response of PAO1 following infection in a bladder epithelial cell line (28). Their study found approximately 1200 DEGs in their intracellular condition, with T3SS and pyoverdine biosynthesis up-regulated. While the number of DEGs between the two studies is similar, we did not observe the same pathway up-regulation in the conditions tested. This could be due to the different conditions between the two studies since their study used cultured epithelial cells infected for two hours with PAO1, which then were sorted for intracellularly infected cells and subsequently enriched for the PAO1 transcriptome.

Studies have shown that the E-D pathway generates NADPH (26, 29, 30). NADPH serves as a crucial cofactor for enzymes involved in oxidative stress responses, including catalase, thiol reductase, and superoxide dismutase (31–33). However, our results show minimal changes in gene expression following the disruption of the E-D pathway. Even within the E-D pathway genes, aside from the deleted gene in the mutants, the basal expression of other E-D genes remains unaltered. This suggests that the *hexR* repressor, which represses *zwf-pgl* when carbohydrate inducers are absent (25), does not significantly affect their expression in the conditions used for these experiments. We compared our findings with a study in PA14 that investigated the transcriptome under flow conditions (34). Follow-up studies found that media contains micromolar amounts of hydrogen peroxide (H_2_O_2_) which is neutralized by cellular catalase (35). Therefore, the bacteria exposed to high flow rates will experience oxidative stress due to the replenishment of the H_2_O_2_ which exceeds inactivation rates (35). Reviewing the initial transcriptomic data for bacteria experiencing high flow vs no flow, the authors identified 38 genes up-regulated over 4-fold after four hours of flow exposure (34). From our mouse instilled data, we identified 38 genes that were up-regulated in one of the E-D mutants as compared to parental PA14. Four of these genes overlapped in the two data set represent 5% of differentially regulated genes (**Supplementary Figure 5**). The majority of these up-regulated are regulated by Zwf since the Δ*zwf* mutant had the majority of these differences. Despite some similarities between differential gene expression of the E-D mutants and bacteria experiencing oxidative stress, the small overlap suggest a limited response to oxidative stress in the E-D mutants under our experimental conditions.

To date, the regulatory effect of the E-D pathway on other genes in *P. aeruginosa* remains unknown. When comparing the mutants of the E-D pathway with the parental PA14 in each condition, there were surprisingly few DEGs. The DEGs were primarily related to energy metabolism, and their expression patterns did not show consistency across mutants of coregulated genes. Each of the E-D mutants had between 1 and 36 DEGs in any given condition, and most of these were associated with the *in-vivo* bladder-instilled condition. Many of these genes were unique to a specific strain and condition; only 18 DEGs were shared among more than one of the E-D mutants, compared to the rest of DEGs (96 genes) unique to individual strain. When individual mutant strains were assessed for DEGs between different conditions, only 12 DEGs were identified in more than one condition for all of the mutants analyzed. The rest of the DEGs were unique to the strain and condition. While there was no consensus transcriptional signal among the E-D mutants, future studies extending the exposure time may identify differential responses downstream of the E-D pathway.

Overall, this study found that deletion of individual genes of the E-D pathway contributed little to *P. aeruginosa* infection in a murine model of CAUTI. Comparing our results with other in vivo studies suggests that the E-D pathway plays different roles in the pathogenesis of various pathogens. In a gastritis model using inbred mice, *H. pylori edd* mutants displayed a 5-log decrease in the gastric mucosa than the wild-type (15). Experiments with *V. cholerae* showed that an *edd* mutant reduced the expression of *ctxA, tcpA*, and *toxT* in vitro and had a 20-fold defect in fluid accumulation in a rabbit ileal loop model (14). In *E. coli* MG1655, an *edd* mutant had a *>* 3-log colonization defect in the gastrointestinal tract (16). Another study using K-12 and F-18 strains reported the *eda* mutant unable to colonize the gut (17). Thus, the E-D pathway contributes to the pathogenesis of gastrointestinal pathogens, such as *H. pylori, V. cholerae*, and *E. coli*. In contrast, E-D mutants significantly increased virulence and decreased survival in a *S. pneumoniae* otitis chinchilla model, leading to decreased host survival (13). Lastly, competition experiments of uropathogenic *E. coli* (UPEC) in a murine UTI model found the *edd* mutant colonized the bladder and kidneys similarly to the wild-type (18). Here, we report similar findings for E-D mutants of *P. aeruginosa* in a CAUTI model. However, several distinctions exist between the two studies. First, the assessment of fitness was conducted after 48 hours post-infection using inbred mice, while our study lasted for two weeks using outbred CF-1 mice. Second, UPEC contains a functional glycolysis pathway in addition to the E-D pathway, and while the E-D mutant can use glucose for in vitro growth, the glycolysis mutant cannot. Therefore, the metabolic capabilities of the two species are different. Third, the study involving UPEC also studied the fitness of other central carbon metabolism pathways in vivo and identified colonization defects in the gluconeogenesis and TCA mutants. This observation could be attributed to the fact that mouse urine contains a higher concentration of proteins compared to glucose (36, 37). Therefore, the significance of the E-D pathway appears to vary across different strains and infection sites. Notably, the E-D pathway may hold particular relevance in UTIs among diabetic individuals, as these infections are notably more prevalent in this population (38). This heightened susceptibility was demonstrated in a diabetic mouse model, where increased vulnerability to UTIs caused by *E. coli, Klebsiella pneumoniae*, and *Enterococcus faecalis* was observed (39). While our CAUTI model did not reveal a significant impact of the E-D pathway on virulence, it may play an important role in a diabetic mouse model.

## Materials and Methods

### Collection of urine

Human urine was collected from healthy adult volunteers after a written informed consent was obtained in accordance with the Institutional Review Board at the University of Maryland, College Park. Human urine was sterile filtered through a 0.2 *µ*m filter, pooled, and stored at -80°C until further use. Mouse urine was collected from anesthetized mice by transurethral insertion of sterile catheter tubing prior to infection. Mouse urine was pooled, sterile filtered through a 0.2 *µ*m filter and stored at -80°C until further use.

### Strains and growth conditions

*P. aeruginosa* PA14 strain was used as a parental wild type strain (40) (**Supp. Table 3**). In-frame deletion mutants were made by PCR amplification of 1 kb regions upstream and downstream of the gene using the indicated primers (**Supp. Table 4**) and ligated into pCR-Blunt (Fisher Scientific). Each of the genes were PCR amplified using the indicated primers (**Supp. Table 4**) and ligated into pCR-Blunt. All fragments in pCR-Blunt were sequence verified. The two 1 kb fragments were sub-cloned into pEX-Gn (41). The deletion construct was introduced into parental PA14 by conjugation using *E. coli* HB101 pRK2013 helper strain (42). In-frame deletion mutants were selected by gentamicin followed by sucrose counter-selection. Mutants were confirmed by colony PCR and sequencing of the PCR product. Complementation plasmids were generated by restriction digest of the gene from pCR-Blunt and cloning into pMMB vector. Complementing plasmids were introduced into the corresponding deletion mutant via conjugation. All strains were grown from a single colony in Lysogeny broth at 37°C shaking unless otherwise stated. For growth on minimal media with defined carbon sources, bacteria were grown overnight in 1x MOPS (43) with 20 mM citrate with 50 *µ*g/mL carbenicillin to select for the plasmid. Next day, the cultures were serial 10-fold diluted in 1x MOPS without carbon source. Five *µ*L of each dilution was spotted on agar plates with 1x MOPS with 20 mM of the indicated carbon source (glucose, gluconate, or citrate) and 50 *µ*g/mL carbenicillin. For growth on complex media, bacteria were grown overnight in LB with 50 *µ*g/mL carbenicillin. Bacteria were serial 10-fold diluted in LB and spotted on the plates with either 75% human urine or LB with 0.5 M urea. Where indicated, 1 mM IPTG was added to induce expression of the indicated gene from the plasmid.

### Murine CAUTI model

The CAUTI model was used as previously described (7, 8). CF-1 female mice (19-21 g) from Charles River Laboratory were group-housed and acclimated for a week. All procedures followed University of Maryland’s IACUC-approved protocol (R-21-30). Mice were anesthetized with ketamine (144 mg/kg) and xylazine (14.4 mg/kg). PE50 catheters were cut into 5 cm guides or 0.4 cm implants and treated with ethyl acetate. Catheters were sterilized under UV light for 10 minutes. Periurethral regions were decontaminated with 70% isopropanol. A 0.4 cm implant was loaded onto a sterile 10 cm metal stylet, inserted into the bladder, and deposited. Urine was voided through the guide. Bacteria cultured to an OD_600_ of 1.0, centrifuged, and resuspended in PBS-tryptone (1.0 × 10^9^ CFU/mL). A 1-mL syringe with a 23-gauge needle was used to inject 35 *µ*l of the inoculum into the bladder, totaling approximately 3.5×10^7^ CFU. Mice were monitored for 14 days and euthanized if showing signs of morbidity or at the end of the 14 days. During dissection, peritoneal cavities were opened, bladders photographed. Implants and organs were extracted, homogenized in sterile PBS-T, and plated on LB agar to count CFUs. Mice without implants were excluded.

### Conditions for RNA-sequencing

For in vitro conditions, strains were grown in LB to ∼1.0 OD_600_. To measure conditions in LB, 1 mL was spun down and stored in RNAlater. The rest of the culture was resuspended in either PBS 1% tryptone (PBS-T) or sterile mouse urine to a concentration of 1.0 × 10^9^ CFU/mL, incubated at 37°C shaking for 2 hours, and then spun down and stored in RNAlater. All samples were performed in triplicate and stored at -80°C until further processing. For in vivo conditions, strains were grown in resuspended in PBS-T to a concentration of 1.0×10^9^ CFU/mL and infected following the CAUTI protocol mentioned above, but the assembly was left in place after delivery of the inoculum for 1 hour of bladder instillation while mice were anesthetized. The tube was cut close to the periurethral area and a slight pressure was applied on the abdomen to collect contents of the bladder in RNAlater. All samples stored in RNAlater were immediately stored at -80 °C.

### RNA sequencing and determining gene expression

RNAlater was removed by centrifugation for 1 minute at 12,000 g. Total RNA was isolated using RNAzol (Molecular Research Center). RNA purity was tested by PCR amplification with My-Taq Red HS polymerase mix and primers for *clpX* to ensure no bands were generated. The total RNA was depleted of mouse and *P. aeruginosa* ribosomal RNA using Illumina total RNA library preparation kit with Ribo-Zero plus and *P. aeruginosa* specific oPools (IDT Technologies). The libraries were sequenced on Illumina NextSeq 1000 for 50 bp paired read resulting in 8.1 to 14.7 million reads. Trimming was performed using Trimmomatic (44) with a sliding window and minimum length of 36 nucleotides. Reads were aligned to the PA14 reference (RefSeq GCF 000404265.1) and mouse reference genome (RefSeq GRCm38.p6, release 100) using HISAT2 (45) (v2.2.1). HTSeq (46) was used to count genes per sample. Discordant reads were not included in gene counts. Data normalization and differential expression analysis was performed in R (v4.2.2) via DESeq2 (19) (v1.38.3). These deletion mutant strains were compared to parental PA14 grown in each medium condition. PA14 was then compared between medium conditions. The entire dataset is shown in **Supp. Table 5**. Sequencing data is deposited in the National Library of Medicine BioProject database under BioProject ID PRJNA1033684.

## Acknowledgements

We thank the members of the Lee lab for reviewing the manuscript. We acknowledge funding from the NIH R01AI110740 and R01AI142400 to V.T.L. We thank April Hussey and the Brain & Behavior Institute – Advanced Genomic Technologies Core (BBI-AGTC), which is supported by the BBI and the University of Maryland, College Park.

## Figure captions

**Supplemental figure 1:** E-D pathway genes are dispensable for growth on citrate as a sole carbon source. Indicated strains with specified plasmid were serial 10-fold diluted and 5 *µ*L were spotted on the plates with specified carbon source and IPTG concentrations.

**Supplemental figure 2:** Assessment of E-D pathway genes for growth on LB supplemented with 0.5 M urea. Indicated strains with specified plasmid were serial 10-fold diluted and 5 *µ*L were spotted on plates with LB containing 0.5 M urea and indicated IPTG concentrations.

**Supplemental figure 3:** Principal component analysis of RNA-seq samples. Data points are color-coded to represent different strains indicated in the legend, and their shapes correspond to the media used: plus signs represent samples from PBS-T, circles represent LB, triangles represent bladder instilled samples, and squares represent samples obtained from mouse urine.

**Supplemental figure 4:** Venn diagram of the number of four-fold differentially expressed genes in PA14 in each indicated condition compared against PBS-T which were **A)** down-regulated and **B)** up-regulated. There are a total of 5,888 genes analyzed.

**Supplemental figure 5:** Venn diagram showing the comparison of four-fold up-regulated genes for this study (purple) and the study of PA14 experiencing flow (yellow) based on data from (34). **A)** The sum of all differential genes between individual E-D mutants as compared to PA14 when instilled into the mouse bladder. Differentially regulated genes from bacteria instilled in the mouse bladder between **B)** Δ*zwf* mutant vs PA14 and **B)** the Δ*pgl* mutant vs PA14.

